# Length measurement of cholinergic innervation in the tunica albuginea of rat testis

**DOI:** 10.1101/2023.10.16.561716

**Authors:** Luis Santamaría, Ildefonso Ingelmo

**Affiliations:** Department of Anatomy, Histology, and Neurosciences. School of Medicine, UAM. Madrid, Spain; Department of Anesthesiology, Hospital Ramón y Cajal. Madrid, Spain

**Keywords:** Testicular albuginea, Cholinergic innervation, Stereology, Vertical sections, Anisotropy, Fiber length estimation

## Abstract

The testicular albuginea is not a simple receptacle that contains the testis parenchyma. It is a biologically active structure involved, among other things, in the progression of sperm from the seminiferous tubules to the rete testis. This function, attributed to the albuginea myofibroblasts’ contractile properties, is mediated by its vegetative innervation. Therefore, the quantification of innervation seems relevant to deepen the knowledge of the testicular envelope’s physiology. The albuginea contains cholinergic nerve fibers that are not very evident and with a marked directional arrangement (anisotropy). These structural characteristics require that the quantification of the fibers’ length be carried out using stereological measurements, which ensure the absence of bias in the process. The authors have developed stereological tools usually carried out on vertical VUR (vertical uniform random) sections in the present work. This approach’s novelty is that the albuginea is treated "in toto" as if it were a thick section (slice) on which a cycloid grid is applied through different optical planes. Thus, on a pilot study of three specimens, cholinergic nerve fibers’ length density was estimated, as well as the estimates of the coefficient of error of measurements and the biological variability of the sample.

Furthermore, after estimating the albuginea volume, the mean of the fibers’ absolute length can be obtained. This stereological approach can be used after visualization of the immunostained nerve fibers for various markers, as long as the reagents’ penetration into the tissue is ensured. Moreover, it can be used in other biological structures, such as the intestine of small animals, which can also be processed as whole mounts and treated as thick VUR sections.

## 1 Introduction

### 1.1 Structure of the Albuginea

The testicular albuginea is a biologically active structure involved in the progression of sperm from the seminiferous tubules to the rete testis. This function is mediated by its vegetative innervation. Therefore, the quantification of innervation seems relevant to deepen the knowledge of the testicular envelope’s physiology. The albuginea contains cholinergic nerve fibers with a marked directional arrangement (anisotropy). These lineal and directional characteristics require that the quantification of the fiber length be performed using stereological techniques that ensure the absence of bias in the procedure.

This chapter will be structured as follows:

An Introduction containing: A brief description of the anatomical and histological characteristics of testicular albuginea; a panoramic vision of the vascularization, innervation, and physiology of the albuginea, and a general approach to the quantitative study of the innervation of the testicular albuginea.

In the Material and Methods section the methodology followed to estimate the length of cholinergic fibers will be described in detail, including procedures for preparing the albuginea as a whole mount section, along with the rationale for why the albuginea In Toto can be considered a vertical uniform random section. The estimation of the anisotropy of the orientation of the cholinergic plexuses of the albuginea will also be addressed in this section. The methodology for the estimation of the relative and absolute length of the nerve fibers and the calculation of the Coefficient of Error (CE) will be described, together with the levels of contribution to the variance observed in the estimation of L_V_ of the cholinergic fibers of the albuginea. Finally a section of Conclusions is included.

The tunica albuginea surrounds the testis and comprises three layers: the visceral portion of the tunica vaginalis, the intermediate portion, and the tunica vasculosa, a thin layer of loose connective tissue [1]. The vasculosa, in most species, sends a series of connective septa that compartmentalize the testicular parenchyma into lobules. These connective trabeculae contain blood vessels, lymphatics, and nerves [1]. The albuginea completely envelops the testis except for the posterosuperior or mediastinal portion, which presents a solution of continuity occupied by the rete testis, thus allowing the connection of the intratesticular spermatic pathway with the efferent cones of the head of the epididymis [1].

To summarize, the albuginea has three compartments:

a. The visceral portion of the tunica vaginalis, i.e., submesothelial supporting layer, lined by the vaginal mesothelium [2, 3].
b. The intermediate portion, formed by dense connective tissue, with bundles of collagen fibers that are arranged in parallel and intermingled with numerous fibroblasts [1] (Fig. 1A).
c. The tunica vasculosa is formed by loose connective tissue, including small blood vessels [1]. In any case, the tunica vasculosa could also be considered a testicular parenchyma tissue, associated with the seminiferous tubes and the interstitial testicular tissue and immediately adjacent to the albuginea [1, 3].

**Figure 1.**
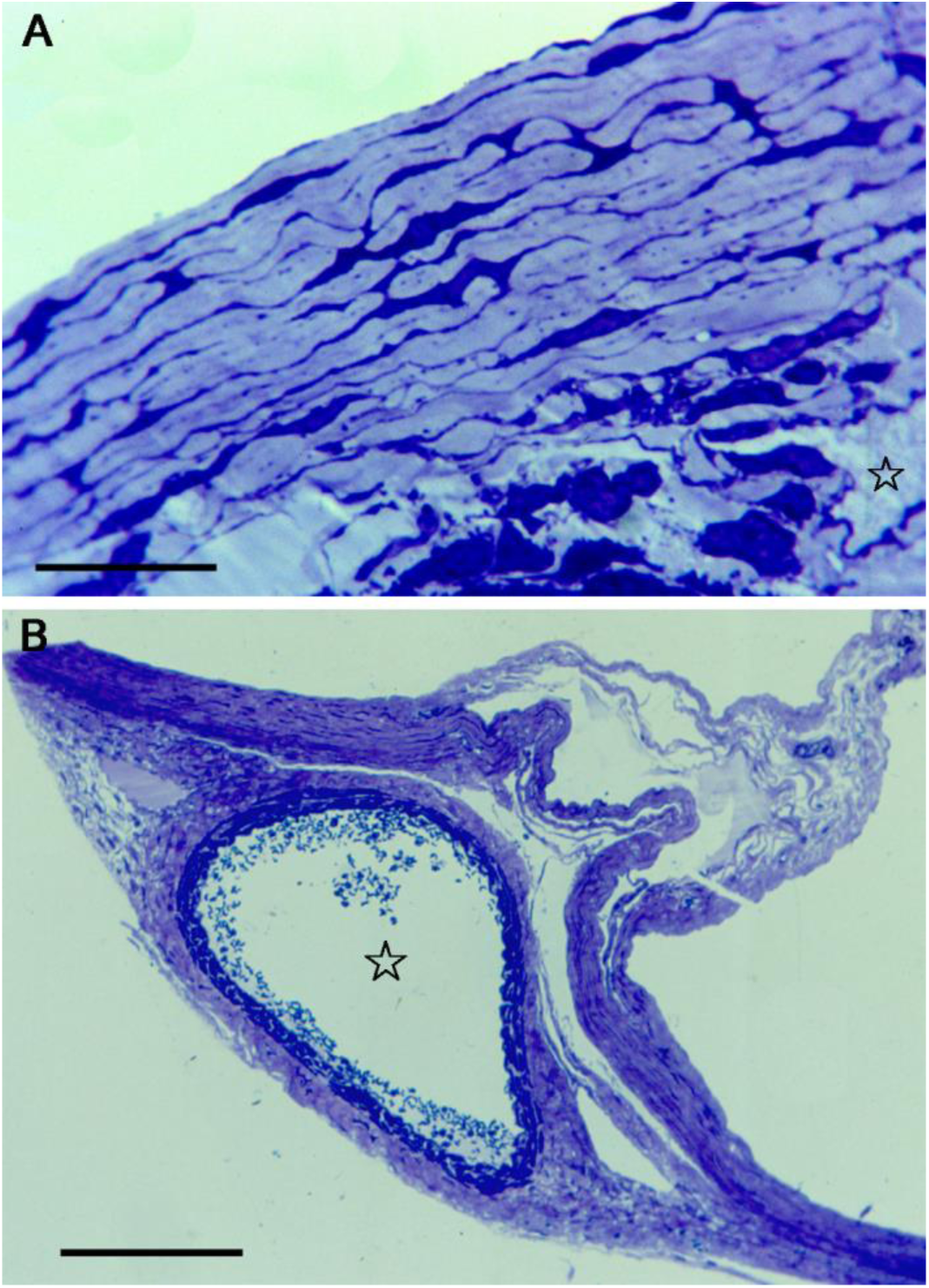
Rat testes were perfused using 10% paraformaldehyde in phosphate-buffered saline. The albuginea was dissected and cut into fragments, which were further fixed in the same fixative for 2 h at 4°C, postfixed in 2% phosphate-buffered osmium tetroxide, dehydrated in ethanol, and embedded in Epon-812. Semithin sections were stained with toluidine blue. (A) Semithin section of rat albuginea; at the top of the image, abundant branched fibroblasts stained by toluidine blue are seen surrounded by a slightly stained stroma. In the lower part of the image is observed a fragment of the tunica vasculosa, with a loose stroma and some blood vessels (star), the scale bar represents 30 µm. (B) Image of the albuginea near the upper pole of the testis, at a lower magnification than in A. In the center of the image, there is a cross-section of the testicular artery (star), the scale bar represents 400 µm.

### 1.2 Vascularization of the Albuginea

In the rat, the testicular artery penetrates at the upper pole level of the testicle and runs as part of the tunica vasculosa (Fig. 1B), along the entire length of the posterior border of the testicleIt has abundant collaterals distributed by the interlobular conjunctive septa, vascularizing the entire testicular parenchyma [1]; however, paradoxically, the tunica albuginea contains few blood vessels [4].

### 1.3 Innervation of the Albuginea

Most species’ tunica albuginea has some small nerve bundles, mostly located near the testicular mediastinum and isolated nerve endings that are irregularly distributed throughout its area [5]. However, according to various authors, the highest proportion of nerve fibers is arranged with blood vessels of the tunica vasculosa [5].

Specifically, the human tunica vasculosa contains unmyelinated nerve fibers, which reach the testicular parenchyma through the interlobular connective septa [6].

Less frequently, bundles of nerve fibers consisting of relatively long axons surrounded by Schwann cells are seen associated with bundles of collagen fibers of the testicular mediastinum. These fascicles and their internal architecture resemble the terminal portion of the extension receptors found in the cat’s articular capsule [7, 8]. Besides, Kreutz W and others [7, 9] have found several types of terminal nerve corpuscles in the tunica albuginea of the man. Kumazawa T [10] verified by stimulating the superior spermatic nerve on the visceral vaginal tunic of the testis and epididymis that polymodal receptors are located in the surface’s close vicinity of the visceral tunica vaginalis.

Relatively thick nerve fibers, evidenced by Bielschowsky’s method, have been found in the visceral vaginal tunic ending in varicosities and free terminals in the tunic’s connective tissue [11]. Multiple receptors have also been found along the surface of the testicular vessels [11]. At present, three different types of testicular receptors are known, depending on the different response patterns to mechanical stimuli applied to the scrotal skin [10, 12].

The testicular albuginea’s efferent innervation is not fully understood [1], although it is well known that the testis receives fibers from the lumbar sympathetic chain [13]. The visceral tunica vaginalis receives efferent nerve fibers from the superior spermatic nerve plexus or nerves related to the testicular artery, innervating the testis’ blood vessels [5, 14-16]. This nerve plexus also participates in afferent innervation by transmitting testicular pain [17, 18]. Some smooth muscle fibers present in the albuginea may be innervated by these effector fibers [1].

Encapsulated nerve endings and free nerve endings have been found in the tunica vaginalis of the cat, dog, and bull [17, 19] as well as in the human testis [14, 20].

Corona GL [17] found Paccini corpuscles and interspersed corpuscles in the testicular vaginal tunic. Kreutz W [9] considers that the encapsulated nerve endings of the testicular tunica albuginea have some similarity to Meissner’s corpuscles and genital corpuscles, but not to Paccini’s corpuscles; besides, he describes a type of nerve ending surrounded by a capsule of connective tissue, as well as another variety of nerve ending contacting with collagen fibers of the tunica albuginea [20].

The modifications that the vaginal undergoes during testis movement may stimulate the Paccini corpuscles of the testicular tunica vaginalis [20]. Also, in the testicular tunic, there are sensitive endings that respond to painful stimuli, and these endings are also located in the wall of the blood vessels; therefore, sustained testicular ischemia can be the cause of scrotal pain [21].

Substance P nerve endings are found in the testicular albuginea and interstitial connective tissue in guinea pigs [22, 23]. In the testis of cats and guinea pigs, small VIPergic varicose endings are distributed in a disseminated way throughout the stroma and around blood vessels of the albuginea, extending through the interstitial conjunctive septa that are arranged between the efferent cones [24]. Peptidergic nerves have not been visualized in the rat [24].

Some findings of histochemical features of cholinergic innervation of rat albuginea were described in a study of Reoyo et al., [25] and could be summarized as follows: The tunica adventitia of the testicular artery that runs across the albuginea from the superior to the inferior testicular poles was seen to contain a rich plexus of acetylcholinesterase-positive (AchE) nerve fibers (Fig. 2A). In the mediastinum testis, collateral branches of this plexus formed a broad network with fibers ending beneath the rete testis epithelium (Fig. 2B). In its course along the testicular artery, the plexus gave rise to many other branches extending laterally over the testis, although without reaching its ventral portion. In turn, these branches originated another series of fibers running parallel to the longest testicular axis towards either the anterior or the posterior poles of the testis (Fig. 2C,D). All these fibers ended inside the albuginea without contacting the scanty blood vessels present in the stroma. The testicular artery exhibited cholinergic nerve endings, which were more abundant immediately before its entry into the testicular parenchyma. Nerve fibers penetrating the meso-epididymis were not observed.

**Figure 2.**
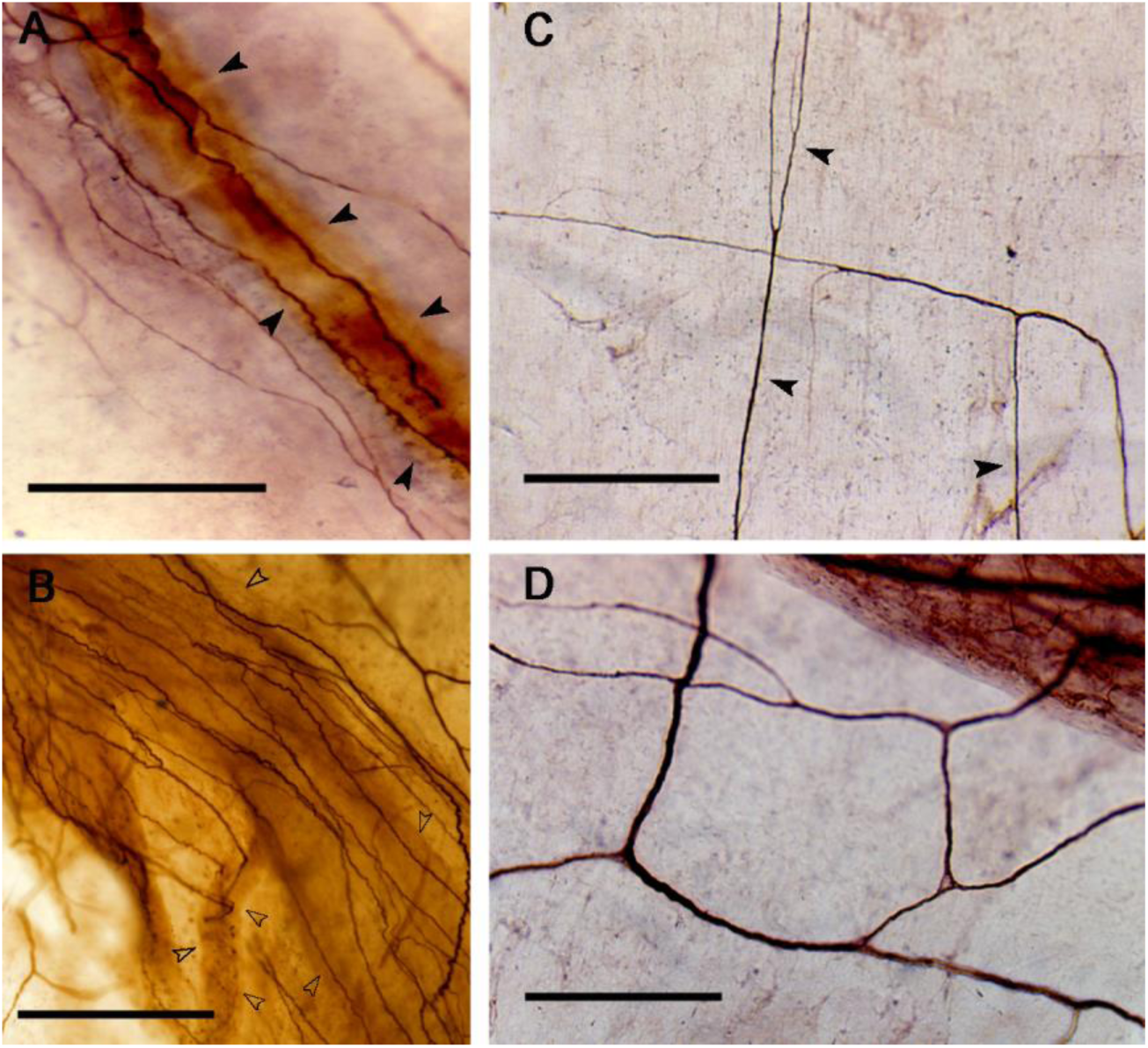
Images from whole mounts of rat testicular albuginea, stained by the Karnovsky-Root method for detecting acetylcholinesterase (AchE) activity in nerve fibers: The tissue was incubated for 40-60 min in a medium containing acetylthiocholine iodide at room temperature. The non-specific activity of cholinesterases was inhibited by adding iso-OMPA, 4 mM, to the medium. After incubation, the tissue was washed in phosphate-buffered saline at pH 7.2, dehydrated in ethanol, and mounted with Depex. (A) AchE positive nerve fibers running parallel to the testicular artery (arrowheads). (B) AchE positive nerve fibers distributed around the rete testis’ ducts are seen by transparency through the albuginea’s thickness (empty arrowheads). (C) The image shows cholinergic fibers that are distributed over great distances through the albuginea. In the picture, those that are located parallel to the long testicular axis are more abundant (arrowheads). (D) Cholinergic fibers of the albuginea. In this field, a more isotropic arrangement in the form of a network is detected. In all images, the scale bars represent 580 µm.

### 1.4 Physiology of the Albuginea

The albuginea has been considered for many years as an inert tissue that surrounds the testicular parenchyma. When albuginea is isolated from testis’ parenchyma, it appears as a saccular structure, weighing about 75 mg in the rat and about twice that in the rabbit [1]. However, albuginea is by no means an inert tissue; on the contrary, it undergoes contractions when it is subjected to the action of acetylcholine and norepinephrine [26-28]; also, it is capable of acting as a semipermeable membrane [1]. Therefore, in addition to protecting the testicular parenchyma, it participates in the regulation of the size of the testicles [1, 29] and probably also participates in some way, still little known, in the exchange between the fluid of the vaginal cavity and the testicular parenchyma, for example in cases of hydrocele: the increase in the intravaginal fluid could cause osmotic changes through the albuginea that modulate the entry and exit of testicular metabolites [1].

The development of the smooth muscle fibers of the albuginea has an essential functional significance [7]. These cells probably participate in changes in intratesticular pressure, facilitating the movement of testicular fluid and sperm [7], which are immobile when they cross the rete testis [30-32]. It does not seem difficult to accept that both the smooth muscle fibers of the albuginea and those of the rete testis can cause extrusion of the contents of the rete testis into the efferent cones.

Acetylcholine and norepinephrine produce a marked contraction of the isolated testicular capsule [33, 34], and this contraction is modified by changing the concentration of the neurotransmitter, obtaining a maximum response with a dose of 1 µg / ml, this maximal contraction occurs approximately three minutes after the addition of the acetylcholine.

Cholinomimetic agents such as carbazole produce a marked contraction of the isolated rat albuginea [26, 33]; however, the contraction response due to a parasympathomimetic drug as pilocarpine is lesser than with carbazole.

The activity of sympathomimetics on isolated albuginea has been also studied epinephrine causes a more significant albuginea contraction than that produced by norepinephrin; however, isoproterenol, which is a beta adrenergic receptor agonist, causes relaxation of albuginea [1, 34]. The sympathetic ganglia’s stimulating agents, such as tetramethylammonium, produce the contraction of the rat’s isolated albuginea [26]. These physiological findings could be related to the relative richness of testicular sympathetic innervation [14] and the supposed absence of parasympathetic innervation cited by some authors [35]. Histamine induces a peculiar contraction of the isolated testicular capsule so that the maximum contraction is followed by immediate relaxation [1]. Other agents with the capacity to exert a contraction in the testicular capsule are oxytocin [36-38] and some prostaglandins [1]. Barium chloride, an intense stimulant of smooth muscle fibers, only produces contraction of the albuginea at extremely high concentrations, which suggests that albuginea has low sensitivity to barium ions, and this could be a consequence of the low number of fibers of smooth muscle [1].

Among the drugs that decrease capsule contractions’ amplitude are: reserpine (anthypertensive drug), atropine (parasympatholytic agent), and a non-steroidal anti-inflammatory drug as indomethacin [34]. In contrast to the rat’s albuginea, that of the rabbit undergoes spontaneous contractions with an interval of 3 to 5 minutes, which shortens approximately 5% of the tissue’s total length [1]. When acetylcholine is administered, a 15% shortening is achieved, and when norepinephrine is added, the contraction reaches 20% [1].

The increase in temperature from 32°C to 38°C, regardless of drugs’ action, produces an intense contraction of the albuginea isolated from the rat, which does not change when greater hyperthermia [1]. The spontaneous contractions of the testicular capsule, are probably related to the transport of immotile sperm.

Also, there are many studies on the stimulation of the testicular capsule by various agents: electrical excitation and norepinephrine [39, 40], F1α prostaglandins [41], hypoxia or anoxia, and epinephrine [42]. The different intervals between stimulation and maximum response could be due to the number of myofibroblasts, differentiation of cytoplasmic filaments, and other plasma membrane receptors [40]. In vitro studies [39] suggest that adrenergic terminal fibers evoke the rat testicular capsule’s contraction. However, the number of adrenergic axons is low compared to the number of myofibroblasts. Thus, the specialized membrane contact of myofibroblasts may be of great importance [40].

For the efficient passage of spermatozoa from the seminiferous tube to the epididymis, the close coordination of a series of factors is necessary, including the ciliary movement of the epithelium of the efferent ducts [43, 44], secretion of seminiferous tubules [45], and the individual contractions of the tubules due to the action of the myoid cells surrounding the tubules [46-51]. Smooth muscle cells of the epididymis act in the complex mechanism of ejaculation and the filling of the tail of the epididymis after sperm emptying [52]. The contraction of the albuginea that surrounds the rete testis could act as a valvular mechanism to regulate the exit of spermatozoa towards the extratesticular spermatic pathway. [52].

The importance of the cholinergic innervation of the testicular albuginea in relation to the mobilization of spermatozoa towards the rete testis is well established.

Therefore, its histological quantification is relevant. Fiber length measurement is the most appropriate estimator since nerve fibers can be considered essentially as linear elements.

### 1.5 Approach to the Quantitative Study of the Innervation of the Testicular Albuginea

Former studies [25] have estimated nerve fiber densities per unit area as a parameter for quantifying the cholinergic innervation of the rat albuginea. These measurements were not carried out by stereological methodology but by applying classical morphometry to evaluate the number of intersections of the cholinergic fiber profiles with a quadrangular lattice. It was observed that the density per unit area of the innervation of the dorsal region was significantly higher in the major axis than in the minor axis of the testis.

Purely qualitative morphological observations detect a remarkable directional structure of the albuginea cholinergic nerve plexuses [53]. Therefore, for an adequate quantification of these fibers, it is necessary to ask about their possible anisotropy. To assess the degree of spatial ordering of the nerve plexuses, Reoyo et al. [25] compared the innervation densities along the major and minor axes of the albuginea considered in two dimensions, but the methods developed did not rigorously address the estimation of possible anisotropy in the orientation of cholinergic fibers.

A correlation between physiological, histochemical, and ultrastructural aspects of both cholinergic and aminergic nerves of rat albuginea, along with the immunohistochemical and ultrastructural analysis of albuginea myofibroblasts, has also been described by some authors [40, 41, 53, 54]. The application of techniques for demonstrating cholinergic and aminergic nerve fibers in whole mounted preparations of the tunica albuginea revealed a greater abundance of nerve fibers than suggested in previous reports [5, 28, 33, 55]. In these reports, the occurrence of cholinergic innervation in the mammalian tunica albuginea was minimized. With the whole-mounted method, a cholinergic innervation as extensive as the adrenergic innervation was demonstrated; this agrees with the results of pharmacologic and functional studies that have placed great emphasis on the cholinergic innervation of the albuginea. The method used provides new information about the distribution of cholinergic nerve fibers in the testicular albuginea.

Hodson N [20] assumed that nerves originated from the hypogastric or pelvic plexuses, accompanying the spermatic artery, branched out at the albuginea level to penetrate the testicular parenchyma and caput epididymis, whereas the nerves accompanying the vas deferens and its vessels only supply the cauda epididymis.

Lamano-Carvahlo et al. [56] have shown that the hypogastric and pelvic decentralization has no noticeable effects on the amount of noradrenergic and peptidergic nerves in the testis and suggested that most aminergic nerve fibers of the testis originate from other sources than the cholinergic nerves. The occurrence of a second source of aminergic innervation agrees with observations of aminergic nerves penetrating in the lower testicular pole by the meso-epididymis of the cauda epididymis.

The ultrastructural pattern of vesicle-containing nerve fibers suggests that they are cholinergic because their 60 nm vesicles lacked the electron-dense granules characteristic of adrenergic nerve vesicles [6, 57]. However, since specific methods for ultrastructural demonstration of neurotransmitters were not used, the possibility that many of the electron-lucent vesicles are adrenergic should not be excluded. The true nature of the neurotransmitters can only be discussed based on the results of the histochemical study.

The occurrence of an extensive plexus of cholinergic fibers among the rete testis channels [53] suggests that such a plexus is related to the contraction of the microfilament rich cells present exclusively at this location. According to Leeson TS and Cookson FB [52] and Leeson TS [29], these cells are smooth muscle cells similar to those reported in the albuginea of several mammalian species, including the dog, the cat [52], the rabbit [58] and man [43]. However, according to their ultrastructural features, the microfilament-rich cells reported [53] were similar to myofibroblasts [40, 41, 54, 59]. In any case, the histochemical demonstration of abundant actin microfilaments in their cytoplasm strongly suggests that they could be contractile cells. The cells in the albuginea’s remaining regions seem to be fibrocytes or inactive fibroblasts with scanty microfilaments and low actin expression. The location of these contractile cells in the rete testis supports the idea that they may be involved in the pumping of semen towards the ductuli efferentes [28, 52]. Other functions of the tunica albuginea, such as the regulation of testicular volume and expulsion of the seminiferous tubule content, are suggested in other mammals, including man [1, 50] but not in the rat, because contractile cells are lacking in the remaining regions of the albuginea. Because true synaptic contacts between cholinergic nerve fibers and contractile cells have not been observed [53], these cells’ contractility is probably regulated by acetylcholine release in their vicinity.

Aminergic fibers are mainly confined to the inferior pole of the albuginea, which precludes the idea that they might be involved in regulating contractile cells. In addition to a local effect, the neurotransmitters released by these fibers could diffuse through the albuginea inside the testicular parenchyma, where they might act on the seminiferous tubules and interstitial cells. In this study [53], mast cells appeared closely related to adrenergic nerve fibers; this agrees with the notion of direct regulation of mast cell secretion by adrenergic nerve fibers as reported in other organs and tissues, including the gastrointestinal tract [60], mucosae [61], the trigeminal ganglion [62] and the sympathetic ganglia [63].

## 2 Material and Methods

### 2.1 Stereological Estimate of Length of Cholinergic Fibers

The rat albuginea, once separated from the testicular tissue, extended onto a plane and manipulated as a whole mount, can be considered as a tissue slice, capable of being oriented and treated as a thick vertical section. Therefore, an approach to the stereological measurement of the rat testicular albuginea cholinergic fibers’ length can be made by applying the length estimation methodology on thick vertical VUR- type sections [64]. First of all, fundamental concepts concerning vertical sections and the generation of test lines with isotropic orientation should be taken into account.

They are essential for estimating linear parameters that, due to their structure, can be anisotropically oriented in space [65].

A practical method suitable for generating isotropic lines on vertically sectioned material was first developed by several authors [66, 67]. An arbitrary reference horizontal plane has to be selected for the object of interest; this reference plane can be chosen as a convenience and can either coincide with a useful plane within the object or be arbitrary. Once the horizontal plane has been chosen, it is then considered as fixed. We can then generate isotropic orientations in the horizontal plane and section the specimen with uniform random position along these orientations. These sections are vertical uniform random (VUR). Remarkably, a VUR section is not an isotropic section in 3D space.

Furthermore, lines of arbitrary orientation in a VUR section do not constitute a set of isotropic lines: a set of lines oriented so that their length density is proportional to sine θ is required [65]. When a cycloid is aligned so that its minor axis is parallel with the vertical direction, it has a direction distribution proportional to sine θ [67, 68].

Combining a plane generated with a VUR protocol and a grid of cycloids oriented in this manner is equivalent to a collection of isotropic uniform random (IUR) lines in 3D space. When using VUR sections, it is crucial to explicitly know the vertical axis direction when the section is finally prepared and mounted on the microscope slide. As indicated before, the testicular albuginea could be considered as a thick section that can be oriented, defining a vertical axis (the major axis of the testicular ellipsoid) (Fig. 3 A), and treated as a VUR section. An approach to the unbiased estimation of length density that uses projections through the thick sections could be employed to measure the length of cholinergic fibers from albuginea. As it is already known [65], the feature length can be estimated by counting the number of intersections between the linear feature and isotropic uniform random (IUR) planes in 3D space. A ’virtual’ IUR surface can be generated in a thick vertical section by projecting a cycloid through the section, if the cycloid’s major axis is parallel with the vertical direction [69]. A vertical uniform random slab of known thickness t, which contains some linear features, is projected onto a cycloidal test system. The intersections between the virtual cycloidal surface and the linear feature in the 3D slab are seen on the projection as intersections between 2D linear elements and the 2D cycloid curve.

**Figure 3.**
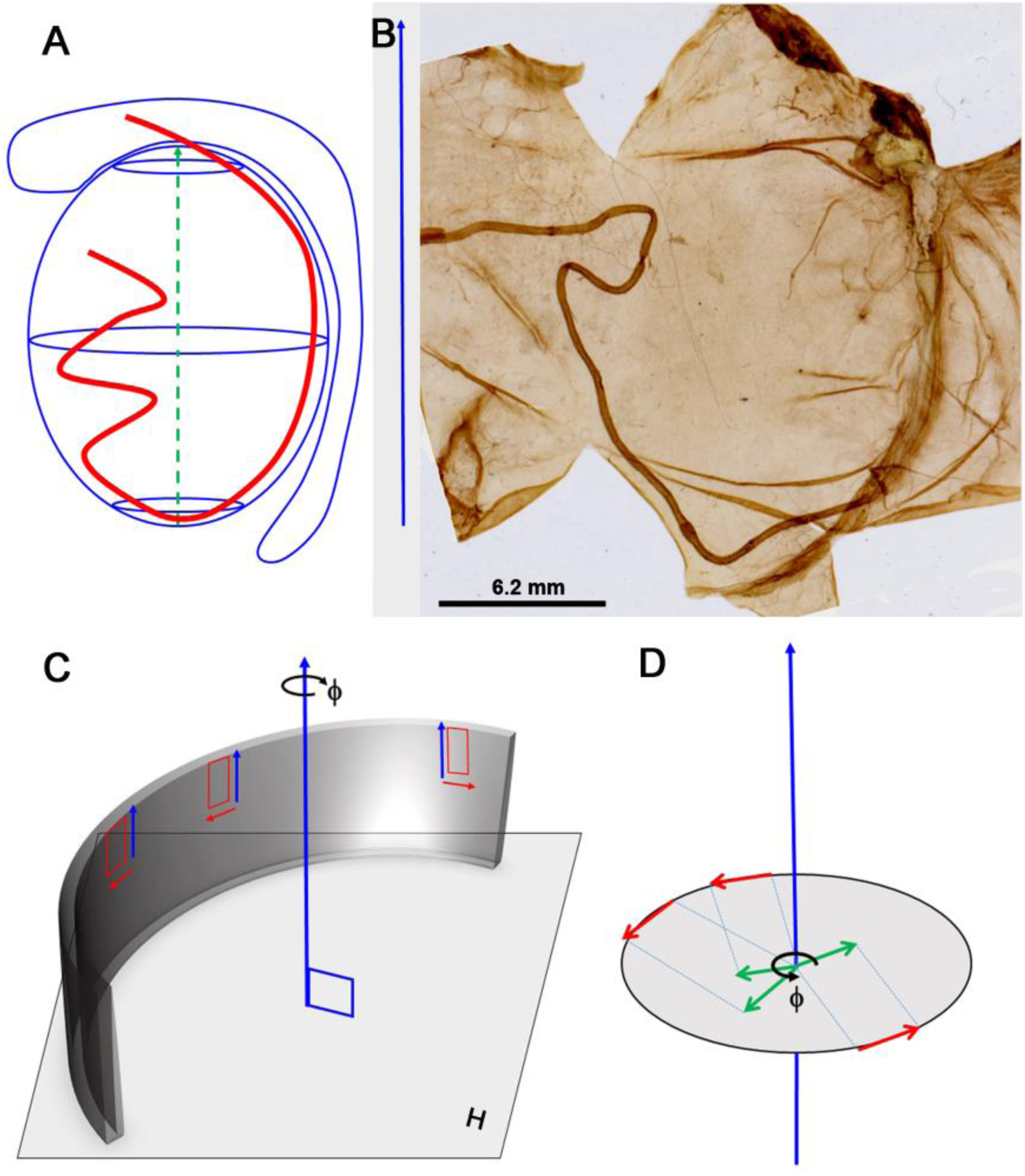
A scheme that indicates the orientation of the albuginea as a vertical thick section VUR (vertical uniform random). (A) Outline of the rat testis, where the testicular artery (in red) is represented penetrating the upper pole of the testis and running towards the lower pole along the dorsal border. Starting from the lower pole, the vessel bends and follows a flexuous path. The major axis of the testicular ellipsoid (dashed green line) is considered the vertical axis (perpendicular to the horizontal plane, i.e. the equatorial plane of the testis). (B) Whole mount of the albuginea, extended in a plane and processed to visualize the cholinergic fibers. The vertical axis is represented by a blue arrow on the side perpendicular to the scale bar. (C) The albuginea is represented as a gray curved surface perpendicular to the horizontal plane H. The vertical axis perpendicular to H is shown. The angle ϕ indicates that the fields contained in the vertical section should be selected according to a uniform random rotation of ϕ between 0 and 360°. The fields that meet this requirement are represented as red rectangles contained in the albuginea: the larger side of each rectangle (blue arrow) is always parallel to the vertical axis, while the smaller side (red arrow) is arranged perpendicular to the vertical axis but with orientations specified by ϕ. (D) A more detailed representation of (C). The horizontal plane is shown as a circle centered by the vertical axis (blue arrow). Around the vertical axis, the directions of the selected fields are represented as green arrows with random uniform rotation (angle ϕ) that translated on the circumference that delimits the horizontal plane (red arrows) show the uniform random orientation of the selected fields on the surface of the albuginea.

Unlike surface estimation [65], when cycloids are used for length estimation in vertical slabs, the cycloid’s major axis is parallel to the vertical direction.

These results were discovered in 1990 by Gokhale AM [69, 70] initially for thick sections. Cruz-Orive and Howard CV developed the design-based formulae for total curve length in 1991 [71]. Vertical slice techniques have been used to estimate the length of blood vessels [72-74], dendritic trees [75], and, more recently, nephron tubule segments [76]. To treat the rat testicular albuginea as a thick section, we first describe the technique used to separate it from the testicular tissue and spread it in a plane.

### 2.2 Method to Obtain the Rat Albuginea as a Whole Mount

Three male Wistar rats, weighing an average of 250 g were killed by inhalation of carbon dioxide. The rats were bred and manipulated in accordance with the bioethical standards of international organizations (WMA Statement on Animal Use in Biomedical Research), European Union guidelines, and Spanish State and Local regulations for the use of animals in research, and approved by the Ethical and animal Studies Committee of the Autonomous University of Madrid. After euthanizing the animals a longitudinal incision was made in both scrotal pouchs, and the testes and epididymis were removed, sectioning at the level of the lower third of the spermatic cord.

Given the low density of innervation of the albuginea, it is not possible to study it using inclusion and cutting techniques; for this reason, histochemical methods have been used on *in toto* albuginea preparations, as other authors [1, 77] have done for the study of the innervation of the blood vessels. These techniques enable the morphometric and topographic analysis of the nerve fibers.

### 2.3 Preparation of the albuginea

The testis was freed from the paratesticular structures, then, a longitudinal incision (2-3 mm in length) was made over the entire anterior border of the testicle, from the upper to the lower pole. Next, the testicular parenchyma was carefully extracted, avoiding possible lesions of the testicular artery and the tunica vasculosa. Four additional incisions were made obliquely to the tissue’s long axis (two in the upper pole and two in the lower pole) to facilitate the complete extension of the albuginea in one plane. The albuginea, thus isolated, was washed with cold isotonic serum.

The albuginea was then spread on a flat silicon surface where it was fixed, constantly moistening in isotonic saline (Fig. 3B).

Once the albuginea was obtained, a histochemical technique was used to visualize the cholinergic fibers as indicated above. In the present study, the Karnovsky-Roots technique was used; briefly: the tissue was incubated for 40-60 minutes at room temperature in the Karnovsky-Roots medium that uses acetyl-thiocholine iodide as substrate [78-83], modified by EI-Badawi [84]. The incubation process is monitored under the microscope, stopping when the nerve plexuses’ satisfactory staining is observed (approximately 2 hours). The extended albuginea is then mounted on a slide. The preparation is treated with 1% osmium tetroxide. It is washed in distilled water in five changes of 3 minutes each. The non-specific activity of cholinesterases was inhibited by adding iso-OMPA, 4 mM, to the medium [53, 85]. After incubation, the tissue was washed in phosphate-buffered saline at pH 7.2, dehydrated in ethanol, and mounted in a synthetic resin (Depex, Serva, Heidelberg, Germany).

### 2.4 Estimation of the Dimensions of the Albuginea Before the Stereological Analysis of Innervation

The mean thickness and total volume of the albuginea were estimated using an Olympus microscope fitted with a motorized stage controlled by Cast-Grid’s stereological software (Stereology Software Package, Silkeborg, Denmark). This program monitors the XY displacement of the microscope stage. The Z displacement of the stage was measured by a microcator whose software is incorporated into the Cast-Grid program. With the indicated program and knowing the mean thickness of the tissue, it is possible to estimate the total volume of the albuginea: *Valb* = *Salb* ● *t*; where: *Salb* is the area of the extended albuginea (mm^2^), and *t* is the mean thickness expressed in mm; in the animals studied, *t* (mean ± SD) was equal to 0.240 ± 0.38 mm and *Valb*: 8253 ± 3858 mm^3^.

All these measurements were conveniently corrected for the shrinkage that the albuginea tissue undergoes after handling due to staining, dehydration, etc. The shrinkage factor (SF) was calculated by measuring the fresh albuginea’s thickness as soon as the testis was removed, so that *SF* = *tf* / *t*, where *tf* = fresh thickness and *t* = thickness of the shrunken tissue, *SF* = 1.1

### 2.5 Justification of why the Albuginea in Toto can be considered as a VUR Section

Two requirements [65, 86] have to be met to treat an object as a VUR section: 1-Existence of a vertical axis: in the albuginea, the long axis of the testicular ellipsoid has been chosen as the vertical axis, which in the extended *In Toto* albuginea coincides with the short side of the structure (Fig. 3A, B).

2-Possibility of generating isotropic orientations in the horizontal plane: usually, these isotropic orientations are produced by making sections that maintain the vertical axis, but whose orientation in the horizontal plane is made according to an angle uniformly distributed at random between 0 and 360°. In albuginea, this is not possible because no cuts are made in the tissue’s thickness. However, the albuginea in 3D space is a curved surface that encloses a volume centered by the vertical axis (Fig. 3C). If fields are selected from the surface of the albuginea with a uniform random distribution, actually those fields can be considered elements of the surface of the albuginea whose minor axis is isotropically oriented in the horizontal plane since it would form angles randomly distributed between 0 and 360° with the vertical axis (Fig. 3D).

### 2.6 Sampling Protocol and Image Acquisition

The specimens of albuginea mounted on slides and their vertical axis aligned with the slide’s shortest side were examined with the Cast-Grid system. At low magnification (x4), the total area covered by the albuginea was selected, and using the software’s sampling system (meander sampling), an average of 10 fields was chosen with a uniform random distribution throughout the entire tissue extension (Fig. 4). In each selected field, a total of 10 images were acquired at a magnification of x10 (at that magnification, 512 pixels correspond to 655 µm), each one corresponding to an optical plane 20 µm apart from the next, covering a total range of 180 µm, leaving a guard area of 20 µm on the upper and lower surface of the tissue. Each specimen collection of 10 images was stored in jpeg format and processed with Image J software (version 1.48), developed at the US National Institutes of Health and available on the Internet at https://imagej.nih.gov/ij/index.html [87] to build image stacks with a total thickness of 180 µm.

**Figure 4.**
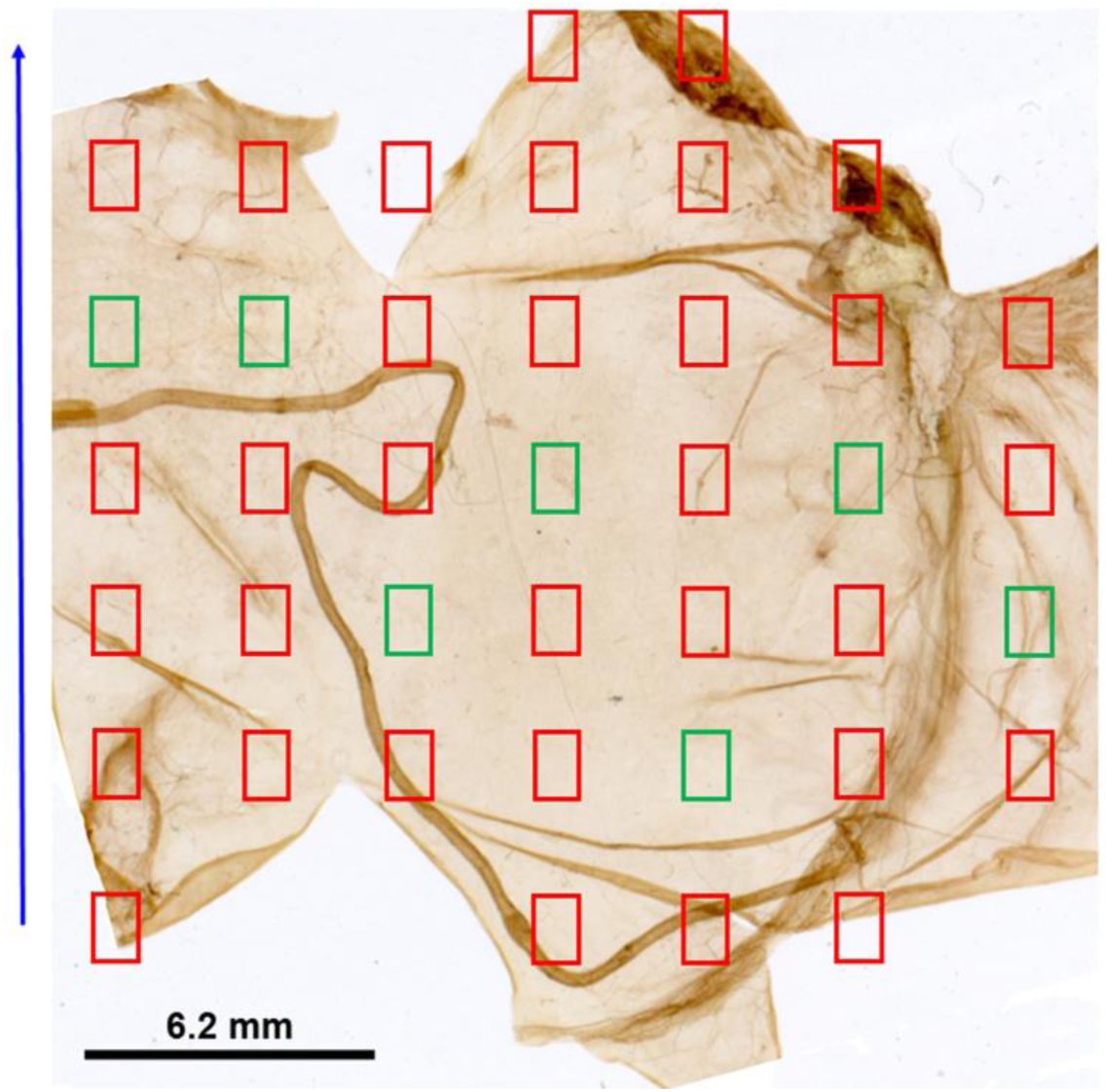
Image showing the entire area of the albuginea extended on a plane. The blue arrow indicates the vertical axis. Using the Cast-Grid software, a total of 40 fields sampled by the "meander sampling" mode (red and green rectangles) has been superimposed on the image. Next, of the selected fields, 10 were uniformly randomly sampled (green rectangles). Given the curved contour of the surface of the albuginea, this selection is equivalent to the one that would be made by rotating each field around the vertical axis, so that the short side of each rectangle is uniformly distributed for 0 ≤ ϕ ≤ 360°.

### 2.7 Investigation about Anisotropy in the Cholinergic Innervation of Albuginea

The findings of some authors [25] suggest that the cholinergic fibers of albuginea show an anisotropic distribution. For a correct stereological approach to the quantification of innervation, it is necessary to discern the presence of the fibers’ spatial orientation, since if a significant anisotropy is observed, the estimation of the length of linear elements in 3D requires the application of suitable tools [65, 86]. Anisotropy analysis is the study of whether spatial pattern differs along different cardinal axes. There are various tools to explore the existence of anisotropy in a structure; for example, the angular correlation, proposed by Simon G [88], is a method of determining anisotropy’s degree in two-dimensional data. This method calculates the correlation between the distance between pairs of points projected onto a vector in a specified direction and the difference in the values associated with those two points. This method is more suitable for analyzing the anisotropic distribution of a population of discrete elements (e.g., cell nuclei) [89]. However, in the present study, when dealing with continuous, linear elements distributed forming a network in 3D space, the estimate of the degree of anisotropy (DA) seems more appropriate: DA is a measure of how highly oriented substructures are within a volume. This anisotropy estimator has been used to analyze the trabecular bone orientation [90] that varies its orientation depending on mechanical load and can become anisotropic. The DA measurement of the cholinergic fibers of the albuginea, considered linear structures with a more or less reticular arrangement, had been carried out using a plugin that runs in Image J. This plugin uses the mean intercept length (MIL) method for determining anisotropy. Briefly, many vectors of the same length originating from a random point within the sample are drawn through the sample. When each vector hits a boundary between foreground and background, an intercept is counted for that vector. The mean intercept length on that vector is then the vector length divided by the number of boundary hits. A cloud of points is built up, where each point represents the vector times its mean intercept length. Fitting an ellipsoid to the point cloud, construction of a material anisotropy tensor and subsequent decomposition results in eigenvalues related to the lengths of the ellipsoid’s axes (as the reciprocal of the semiaxis squared) and eigenvectors giving the orientation of the axes. DA is calculated as 1 - smallest eigenvalue / largest eigenvalue (note that the longest axis has the smallest eigenvalue). New random points with the same vectors are sampled and DA updated with the new MIL counts until either the minimum number of sampling points is reached or the DA variation coefficient below a threshold [91]. The DA was estimated on the image stacks built as indicated above after its binarization, which was also carried out with the Image J software. DA values range between 0 and 1, with 0 corresponding to a complete isotropy situation and 1 to maximum anisotropy.

For more accurate discrimination of DA in real cases, a simulated model of a network of linear elements of isotropic distribution was made as follows:

Three simulations were carried out; in each one of them, a stack of images were built, each consisting of 5 images. Each image was created as follows:

1-For each simulation, a set of pairs of coordinates was obtained, chosen at random from a series of real numbers ≥ 0, according to a Poisson distribution. The average number of coordinate pairs was 110.

2-From each set of coordinates, in each simulation, a network of connections was built using segments following the Delaunay tessellation method [92].

3-Once the connection network was obtained, a series of images rotated 90° respecting to the previous one was generated from each of the tessellated images, so that, finally, for each simulation, a series of 5 images were obtained. Then a stack of five images was made for each of the three simulations. The random selection of the pairs of coordinates to generate the tessellation segments and the rotation of each image of the stack ensures the isotropic distribution of the reticulum that simulates a nerve fibers’ plexus.

The elaboration of the random coordinates and the segment lattices obtained from these coordinates was performed using the PASSaGE software [93], which is a program suitable for pattern analysis and spatial statistics. The construction of the stacks and their subsequent binarization for the estimation of DA was carried out by Image J, similar to that described above for albuginea’s real images.

The DA of the cholinergic fibers of the albuginea is significantly higher (p <0.05) when compared with the values obtained on the isotropic simulated models (Fig. 5a- c), which indicates that the spatial arrangement of the cholinergic fibers is anisotropic. Therefore, the cycloid curves onto thick VUR sections will be the most suitable method for estimating the length density of cholinergic fibers since adequately used; it avoids the bias caused by these fibers’ preferential orientation (61, 62).

**Figure 5.**
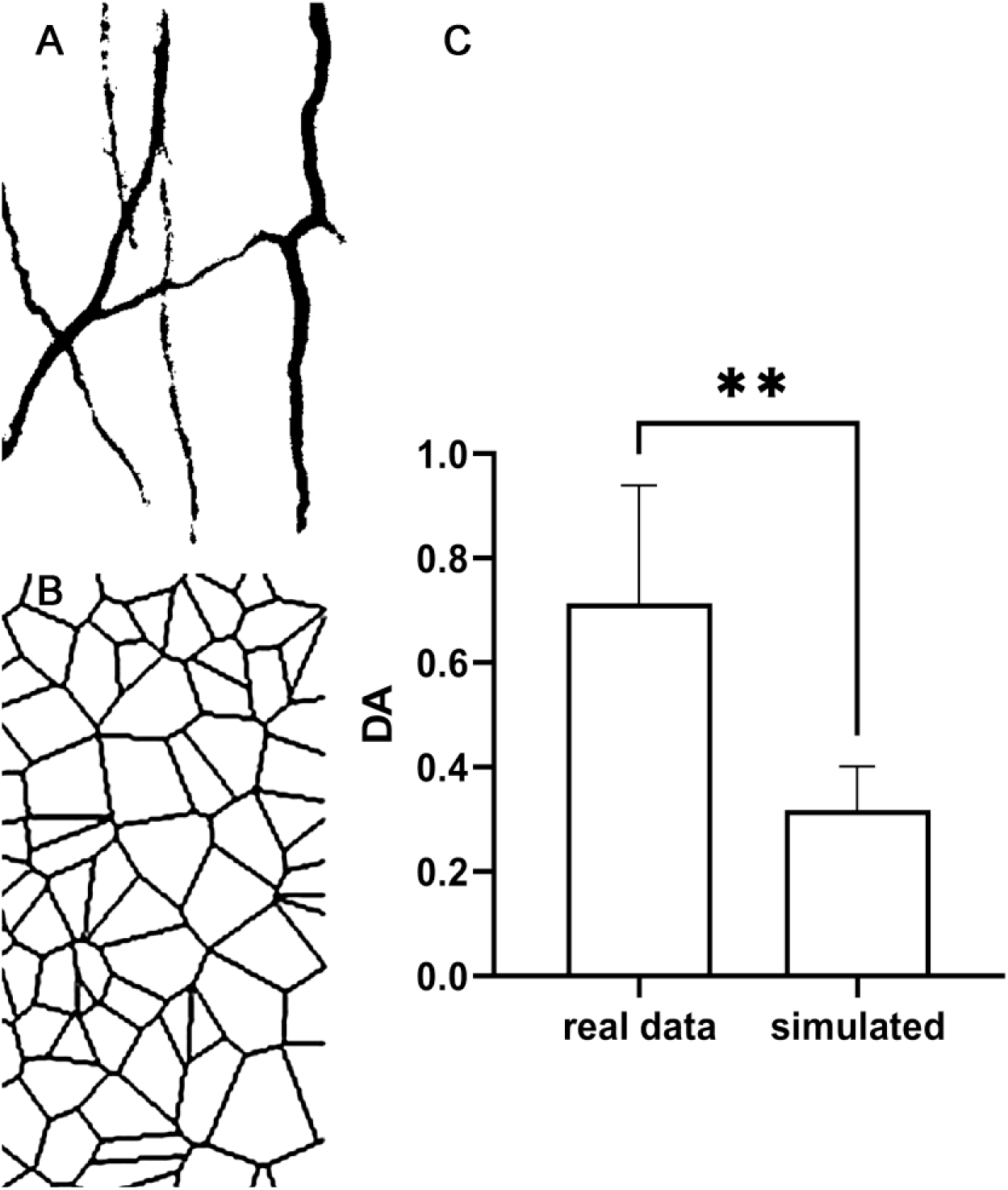
This image shows the evaluation of the degree of anisotropy, carried out prior to the stereological estimation of the length of the cholinergic fibers in the rat albuginea. (A) Binarized image of an optic plane extracted from a stack of images of nerve fibers from albuginea. The degree of anisotropy (DA ± SD) averaged over all the image stacks and over the three cases studied was: 0.71 ± 0.45. (B) Binarized image of a model constructed to simulate an isotropic filament network. In the model, the DA averaged in the same way as for the real cases was: 0.31 ± 0.08. (C) Bar chart in which the average DA is represented both for real data and for the simulated model; error bars indicate SD. After the comparison using the Student’s t test, it was observed that the DA of the real data was significantly higher (p < 0.05) than the DA of the simulated ones. The double asterisk expresses that p = 0.002.

### 2.8 Cholinergic Fiber Length Estimation

The image stacks elaborated as indicated above were employed to estimate the length density of the nerve fibers (L_V_ = fiber length per unit volume of tissue) using an Image J plugin. This plugin consists of a frame of cycloids associated with points that is superimposed on the stack; provided an adequate orientation of the major axis of the cycloid (parallel to the vertical axis of the preparation), it allows count the intersections of the cycloid curves with the profiles of the cholinergic fibers through the different images (optical planes) from the stack (Fig. 6A-C). To optimize the encounter of the curves with the profiles of the nerve fibers, the software allows designing the number of cycloids per frame, as well as the length of the cycloid associated with the point l(p), which in the present study was l(p) = 95 µm.

**Figure 6.**
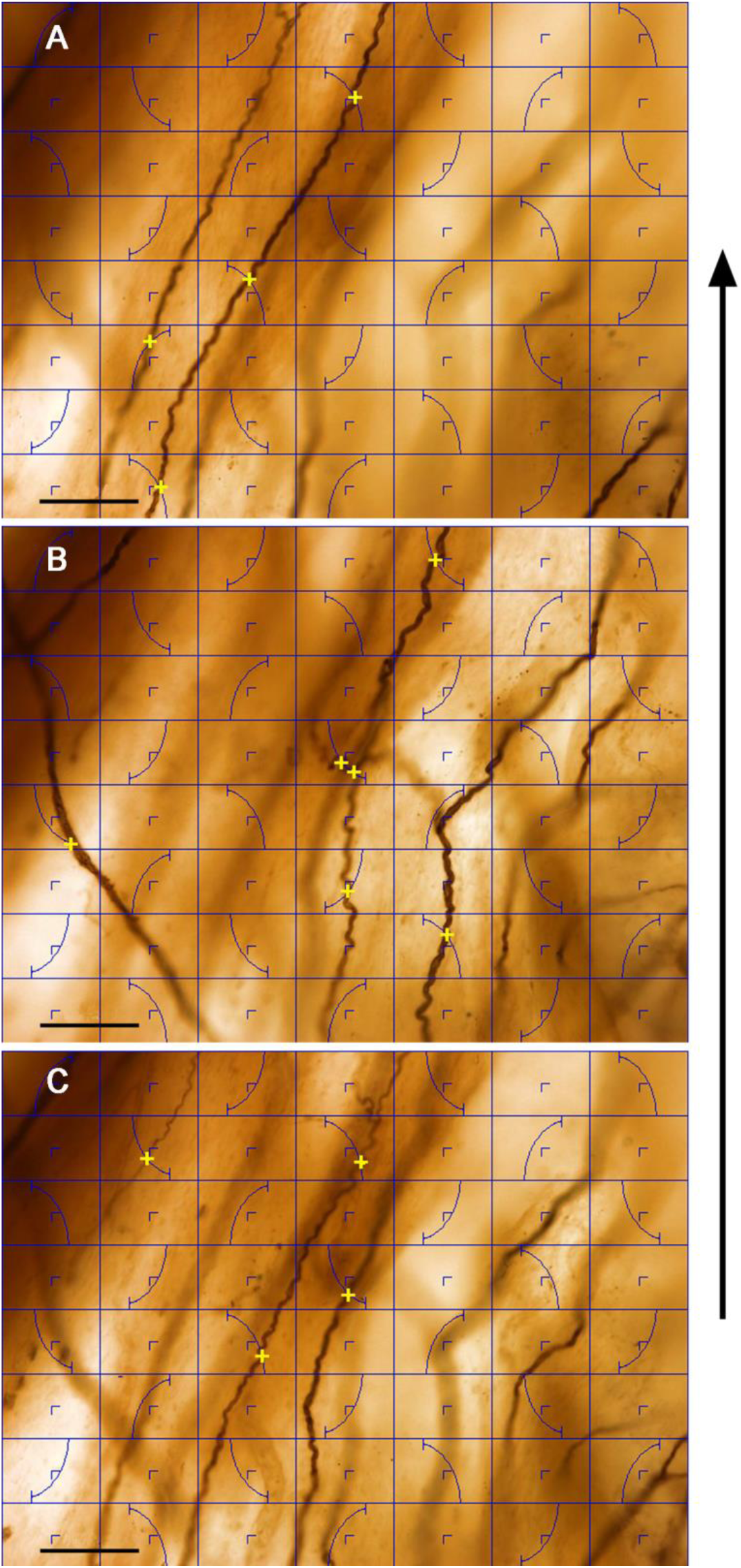
This image shows three consecutive optical planes (A, B, and C) from a stack of images of rat testicular albuginea where cholinergic fibers are visualized. The total thickness of the stack was 180 µm. The three planes shown are 20 µm apart. The Image J software superimposes on the images a cycloid frame whose major axis is parallel to the vertical axis of the images (represented by the black arrow located on the right margin of the composition), this axis retains the same direction of the vertical axis for the entire albuginea (i.e.: The long axis of the testicular ellipsoid). The grid used includes, in addition to the cycloids, a frame of points (represented by small inverted right angles) used to count the number of points (P) falling on the sampled reference space. The length of the cycloid associated with the point, l(p) was equal to 95 µm. On each of the images, the intersections (I) of the cycloids with the profiles of cholinergic fibers focused on each optical plane are counted and marked as yellow crosses. By counting ∑I and ∑P on all the stacks in each case, *L_V_* can be estimated according to the formula indicated in the text. Scale bars from all the images represent 124 µm.

Once the total intersections of the cycloids with the cholinergic profiles have been counted, the following formula is applied to estimate L_V_ [65]:

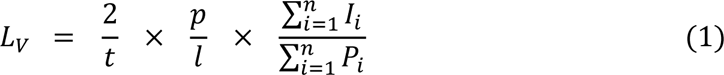

Where:

*t*: thickness of the image stack (180 µm)

*p* / *l*: 1 / l(p).

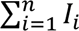: Sum of the number of intersections of the cycloids with the profiles of the fibers.

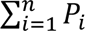: Sum of the number of frame points falling on the sampled reference space (albuginea tissue).

From the *L_V_*, the absolute length (*L*) can be estimated, knowing that: *L* = *L_V_* ● *Valb*; where *Valb* is the total volume of the albuginea calculated as indicated above.

The results of *L_V_* and *L* for three specimens of rat albuginea are indicated in Table 1.

**Table 1.** The results for three specimens of rat testicular albuginea (case) and their corresponding means (mean ± SD) are indicated. Valb (volume occupied by the albuginea expressed in mm3); L_V_ (length density of cholinergic fibers expressed in mm^-2^); L (absolute length of cholinergic fibers expressed in m).

| Case | Valb (mm<sup>3</sup>) | L<sub>v</sub> (mm<sup>-2</sup>) | L (m) |
| --- | --- | --- | --- |
| 1 | 12157 | 6.15 | 75 |
| 2 | 8160 | 2.66 | 22 |
| 3 | 4443 | 5.00 | 21 |
| Mean ± SD | 8253 ± 3858 | 4.53 ± 1.76 | 20.5 ± 3 |

### 2.9 Calculation of the Coefficient of Error (CE) of L_V_

The estimator’s precision for *L_V_* can be calculated by estimating the coefficient of error (CE). Observing the formula used for the estimation of the length density of the cholinergic fibers (see equation 1), it is denoted that *L_V_* was obtained from a ratio between the total of intersections of the positive AchE profiles with the cycloid frame and the total of points falling on the space reference (i.e., the albuginea). Ratio estimators of this type have a small intrinsic bias that rapidly decreases as the number of fields explored increases [94-96], and it can be approximated using the following formula [65]:

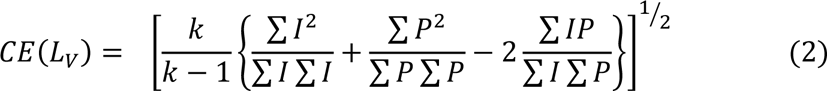

Where:

*k:* is the number of fields measured, and each summation in the formula is over 1 to *k*.

*I:* is the number of intersections between the fibers and the cycloids.

*P:* is the number of points falling on the reference space.

The values of CE and the intermediate steps (operations with I and P) for its estimation are expressed in Table 2.

**Table 2.** The results of the calculation of the coefficient of error (CE) for the estimation of L_V_ are expressed. The calculations of the intermediate stages are indicated for each specimen (Case), using the values of I and P used in equation 2. The averages of CE^2^ and CE are also shown (Mean) for the set of the 3 specimens studied.

| Case | k | $\sum I$ | $\sum P$ | $\sum I^2$ | $\sum P^2$ | $\sum I \sum I$ | $\sum P \sum P$ | $\sum I \sum P$ | $\sum IP$ | CE<sup>2</sup> | CE |
| --- | --- | --- | --- | --- | --- | --- | --- | --- | --- | --- | --- |
| 1 | 10 | 85 | 1600 | 907 | 256000 | 7225 | 2560000 | 136000 | 13600 | 0.03 | 0.16 |
| 2 | 10 | 34 | 1559 | 150 | 244561 | 1156 | 2430481 | 53006 | 5112 | 0.04 | 0.20 |
| 3 | 10 | 66 | 1600 | 520 | 256000 | 4356 | 2560000 | 105600 | 10560 | 0.02 | 0.14 |
| Mean |  |  |  |  |  |  |  |  |  | 0.03 | 0.17 |

### 2.10 Levels of Contribution to the Variance Observed in the Estimation of L_V_ of the Cholinergic Fibers of the Albuginea

The variance of the mean of a sample will have two components: (a) that due to the variation between cases (biological variation) that is inherent to the nature of the specimens studied and (b) the variation introduced by the method used [65].

In this study, and as usual, when estimating the amount of variation introduced by these two components, the quadratic values of the coefficient of variation (CV) and the coefficient of error (CE) are used, according to the formula [65]:

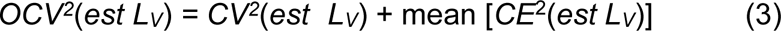

Where:

*OCV^2^*(*est L_V_*) = SD^2^(*est L_V_*) / mean^2^(*est L_V_*): Observed coefficient of variation (squared).

*CV^2^*(*est L_V_*): coefficient of variation (squared) due to the specimens’ biologic variance. mean [*CE*^2^(*est L_V_*)]: average of the coefficient of error (squared) of the method for estimation of L_V_.

Then, we are interested in calculating the biological variance, which informs us about the variability of the length density of the cholinergic fibers due to the characteristics of the animals studied. From equation 3 we know the following terms:

The CV due to the observed variance: *OCV^2^*(*est L_V_*), and the CE due to the method used: mean[*CE*^2^(*est L_V_*)], clearing the term due to the biological variance, it remains:

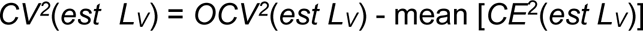

From Tables 1 and 2 we obtain:

*OCV^2^*(*est L_V_*) = SD^2^(*est L_V_*) / mean^2^(*est L_V_*) = 1.76^2^ / 4.53^2^ = 0.15 mean [*CE*^2^(*est L_V_*)] = 0.031

Then:

*CV^2^*(*est L_V_*) = 0.15 – 0.031 = 0.119, then: *CV*(*est L_V_*) = 0.34,if: *OCV*(*est L_V_*) = 0.38.

From this, it follows that the biological variance represents 89% of the observed variance.

## 3 Conclusions

The testicular tunica albuginea is not a simple receptacle that supports and protects the testicular parenchyma [97], but also plays an essential role in the progression of sperm to the spermatic pathways [3], mainly due to the contractile function of its myofibroblasts [27]. Likewise, the albuginea’s innervation’s regulatory role is important, and its quantitative assessment is essential to provide functional data [97]. The quantification of the albuginea’s innervation by applying stereological tools in conventional histological sections is difficult due to the scarcity of innervation and its highly directional distribution [53].

In the present study, it has been shown that the rat testicular albuginea, when processed "*in toto*" (i.e., by a whole-mount technique), can be considered and treated as a thick vertical section, in which it can also be sampled in a number of fields with uniform random distribution around a pre-defined vertical axis (the major axis of the testicular ellipsoid). Thus, appropriate stereological tools have been applied to VUR (vertical uniform random) sections to estimate nerve fibers’ length density. Although other stereological techniques have been recently developed and allow the estimation of the length of linear elements without the need to practice VUR sections (spatial balls method and virtual planes method) [98, 99], in the case of albuginea, the design carried out without the need for physical sections on the material allows the use of cycloids in a direct and straightforward way.

As a preliminary step before applying these stereological tools, we have confirmed the significant anisotropy of the albuginea nerve fibers’ distribution. Although previous studies already suggested the directional component of the distribution of nerve fibers [25], in the present work, the anisotropy of the nerve fibers of the rat testis albuginea has been confirmed through the application of estimators of the degree of anisotropy (DA) [91] and their comparison with models that simulate isotropic networks, and therefore the need to quantify them by applying stereology on VUR sections.

The estimation of the relative length of nerve fibers (length density, i.e., fiber length per unit volume) was performed as described above. Then, the estimate of the absolute length (total length of nerve fiber in the entire albuginea) was obtained by multiplying *L_V_* by the volume occupied by the albuginea, which was previously determined as a result of the product of the area of the albuginea by its average thickness [99]. A dimensional correction of the volume was applied using a factor for the tissue’s shrinkage with fixation, staining, dehydration, and mounting.

Although this work has focused on quantifying cholinergic innervation, this methodology can be applied to whole-mount preparations immunostained for different types of nerve fiber markers (neuropeptides, neurofilaments, etc.). The described technique can be useful to quantify the innervation of other organs or structures that can extend on a plane and have an adequate thickness so that the reagents for immunostaining penetrate and present a degree of transparency that allows the analysis and quantification in different optical planes through the total thickness of the tissue: for example rat intestine that can be sectioned along, extended in one plane, defining the long axis of the intestinal tube as the vertical axis. Then, after adequate immunostaining to visualize the intestinal plexuses, the stereological estimates described in this work could be made.

This work constitutes a pilot study on only three specimens of rat testicular albuginea; several authors have emphasized the interest of this type of study in design-based stereological methods [100]. The small number of cases may explain the existence of a relatively high CE (17%). Increasing n will probably reduce the coefficient of the error to more suitable levels. As already described by various authors [101], to minimize dispersion, it is more effective to increase the sample size than to increase the work intensity on a smaller sample (increase in the number of fields, the counting frame density, etc.). In any case, in the cases analyzed, it is observed that the contribution to variability by biological variance is very high (more than 80% of the observed variance is attributed to biological variability), this factor, as it is intrinsic to the structure of the object of study [65] cannot be reduced and is also indicative of the degree of "natural" heterogeneity of the specimen in question.

